# Transcriptomic characterization of tuberculous sputum reveals a host Warburg effect and microbial cholesterol catabolism

**DOI:** 10.1101/2020.03.09.983163

**Authors:** Rachel PJ Lai, Teresa Cortes, Suzaan Marais, Neesha Rockwood, Melissa L Burke, Acely Garza-Garcia, Stuart Horswell, Anne O’Garra, Douglas B Young, Robert J Wilkinson

## Abstract

The crucial transmission phase of tuberculosis (TB) relies on infectious sputum yet cannot easily be modeled. We applied one-step RNA-Sequencing to sputum from infectious TB patients to investigate the host and microbial environments underlying transmission of *Mycobacterium tuberculosis* (*Mtb*). In such TB sputa, compared to non-TB controls, transcriptional upregulation of inflammatory responses and a metabolic shift towards glycolysis was observed in the host. Amongst all bacterial sequences in the sputum, only less than 1.5% originated from *Mtb* and its abundance is associated with HIV-1 coinfection status. The transcriptome of sputum *Mtb* more closely resembled aerobic replication and was characterized by evidence of cholesterol utilization, zinc deprivation and reduced expression of the virulence-associated PhoP regulon. Our study provides a comprehensive analysis of the transcriptional landscape associated with infectious sputum and demonstrates the feasibility of applying advanced sequencing technology to readily accessible pathological specimens in the study of host-pathogen adaptation.

## Introduction

Concerted efforts over the last two decades have widened availability of therapy for tuberculosis (TB). While this has saved millions of lives, the incidence of disease has declined by only 1.5% annually ^1^. The host-pathogen interaction in TB is complex, thus hindering the development of diagnostic tests and effective new treatments. Studies on TB rely heavily on *in vitro* or *in vivo* experimental models, or blood from TB patients, as lung sampling is invasive. While these approaches provide insights into TB immune responses and the development of tuberculous lesions at a cellular and molecular level, the events following bacterial release from liquefied lung cavities into the airways remain poorly understood.

As TB is spread by aerosol generated mainly through coughing, understanding the physiological state of *Mycobacterium tuberculosis* (*Mtb*) and its interaction with the host in the nasopharyngeal environment may bring insights on new treatment or preventive therapy strategies. Sputum is routinely collected for TB diagnosis and has been proposed as a surrogate for bronchoalveolar lavage for monitoring the transcriptional profiles of *Mycobacterium tuberculosis* (*Mtb*) in patients ^2^. While several studies in the past have characterised the transcriptomes of sputum *Mtb* using microarray and/or targeted quantitative PCR (qPCR), but lacked simultaneous profiling of the host response. We reasoned that a comprehensive RNA sequence-based analysis that yields dual host-pathogen transcriptomes would provide important insight to improve understanding of the biology of *Mtb* transmission and pathogenesis. Technical difficulties and the overwhelming eukaryotic content have limited conventional sequencing approaches either to the host or to a pathogen that has been physically separated or independently enriched, but dual RNA-Seq allows comprehensive and simultaneous survey of gene expression of both the host and the pathogen in one step. To date, there have been increasing success in dual RNA-Seq where the technology was successfully applied to profiled gene expression of *Salmonella enterica* in infected HeLa cells ^3^, *Haemophilus influenzae* colonized primary mucosal epithelium ^4^ and murine Peyer’s patch infected with *Yersinia psedotuberculosis* ^5^. Non-one-step dual RNA-Seq has also been used to study *Mycobacterium paratuberculosis* and *Mycobacterium bovis* Bacillus Calmette-Guerin (BCG) infected cells *in vitro* but with limited success despite separate microbial enrichment ^6,7^. Most recently, dual RNA-Seq on Mtb-infected mice indicated that alveolar and interstitial macrophages utilised different mechanisms to sustain or restrict intracellular *Mtb* growth ^8^. In this study, we applied one-step dual RNA-Seq to sputa collected directly from patients with and without active TB to survey the global transcription profiles of the host and *Mtb*. Transcriptional signature of TB-infected host displayed characteristic of the Warburg effect, while cholesterol catabolism and zinc-deprivation were identified in sputum *Mtb*.

## Results

### Dual RNA-Seq and the host transcriptome

RNA was extracted from 17 sputum samples from South African patients with untreated active TB (9 HIV-uninfected and 8 HIV-infected, referred to as TB-only and TB-HIV, respectively) and 9 samples from persons with respiratory symptoms but no evidence of active TB (referred to as non-TB) (**Table S1**). No physical separation or microbial enrichment was performed to avoid technical error or bias. An average of 1.7×10^8^ reads were generated per sample. Sequence reads were first quality filtered then aligned to the human genome, with unaligned reads extracted for microbiome taxonomy classification and species mapping (**Fig. 1a**). Regardless of HIV-1 status, human reads accounted for an average of 74(±17)% and bacteria for 13(±13)% of all sequenced reads in tuberculous samples (**Fig. 1b**). In contrast, non-TB sputa generated significantly fewer human reads (44±20%, *p*=0.0007) and a non-statistically significant higher number of bacterial reads (24±21%). Unassigned reads may have arisen from incomplete filtering of human sequences and from fungal and unidentified bacterial genomes missing from the database.

**Fig. 1.**
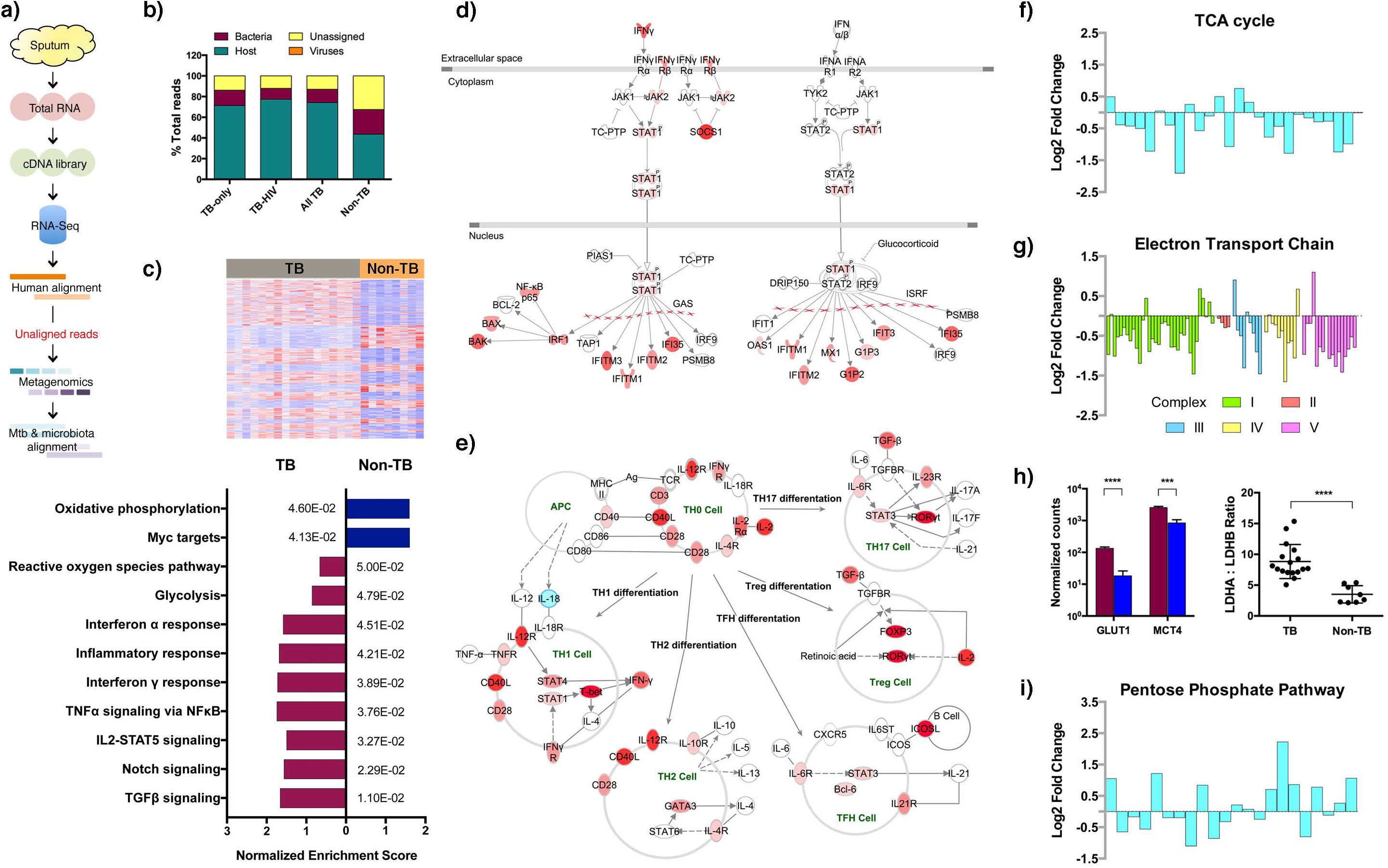
Dual host-pathogen RNA-Seq and the host transcriptome. **a)** Sputum samples were collected from 17 active TB and 9 non-TB respiratory symptomatic patients. Total RNA was extracted and cDNA library generated for ultra-deep RNA-sequencing. Sequence reads were first aligned to the human genome and unmapped reads were extracted for further microbiome metagenomics classification. After identifying the predominant microbiome taxa, reference-based alignment was performed to the top 10 abundant microbiome species as well as to *Mtb*. **b)** Global transcript composition profiles of TB and non-TB sputa were calculated. A reduced percentage of host reads and increased percentage of bacterial reads was recorded in non-TB samples. **c)** Heatmap showing a total of 5843 differentially expressed genes in the host transcriptomes between TB (n=17) and non-TB (n=8) sputa. Gene set enrichment analysis identified 9 pathways that were significantly enriched in TB and 2 in non-TB. The *p-value* of each enriched pathway is listed. **d)** Genes associated with IFNγ and IFNα/β signaling pathways were significantly enriched in TB samples. Red indicates upregulation in TB sputa, compared to non-TB. **e)** Evidence of T cell subset differentiation or recruitment was also observed at the transcriptional level albeit with generally low read counts. Red indicates upregulation and blue downregulation in TB *versus* non-TB sputa. **f) and g)** Metabolic reprogramming was observed in TB sputa, with decreased expression of genes in the TCA cycle and electron transport chain. The log2 fold change of TB sputa compared to non-TB is shown here and indicative of metabolic reprogramming with significant decrease in genes involved in TCA and electron transport chain. Statistical significance of each gene is listed in Supplementary Table S3. **h)** In contrast to decreased oxidative phosphorylation, there was a significant increase of genes associated with glucose uptake and lactate export in TB sputa (red) when compared to non-TB controls (blue). An increased LDHA to LDHB ratio is indicative of conversion of pyruvate to lactate. Statistical significance (p-values) are shown as asterisks: *** padj<0.001 and **** padj<0.0001. **i)** Transcript expression of genes involved in the NADPH production in the pentose phosphate pathway was also significantly higher in TB sputa. A detailed pathway map with the significance of each gene is shown in Supplementary Fig. S3.

We first examined the impact of *Mtb* and HIV-1 infections on the host transcriptome. We identified 21 genes that were differentially expressed in HIV-1 co-infected TB sputa (**Table S2**), including upregulation of T-cell markers such as CD8A/B, LAG3 and CRTAM. This observation was consistent with that from nonhuman primates with TB, in which co-infection with simian immunodeficiency virus significantly induced LAG3 expression ^9^, suggesting that T-cell recruitment to TB sputum is quantitatively and qualitatively affected by HIV-1 co-infection. The presence of *Mtb* had a significant impact on the host transcriptome in the respiratory tract, with total segregation between TB and non-TB samples in Principal Component Analysis (**Supplementary Fig. S1**). One of the non-TB samples (SP321) was a conspicuous outlier and was omitted from further analysis. Comparison between TB sputa (regardless of HIV-1 status) and non-TB controls identified 5843 genes that were differentially expressed (log2FoldChange > ±0.5, *p-adjusted* < 0.05; **Table S3**). Gene set enrichment analysis of these 5843 genes identified 11 significant gene sets, of which 9 were positively enriched in TB sputum and 2 were negatively enriched in non-TB (**Fig. 1c**).

The TB enriched pathways consisted of inflammatory responses mediated by interferon-gamma (IFNγ), tumor necrosis factor alpha (TNF-α) and, to a lesser extent, by type I interferon (IFNα/β) (**Fig. 1d**). The enhanced transcription of these inflammatory mediators is consistent with elevated cytokine concentrations previously reported in TB sputum when compared to pneumonia controls ^10^. Significant transcriptional changes associated with T helper cell activation and differentiation, including T-bet, GATA3, RORγt and FOXP3 transcriptional regulators, were also detected despite lymphocytes typically accounting for less than 1% of the total cellular composition in TB sputum ^10^ (**Fig. 1e**). Expression of IL-18 was significantly downregulated in TB sputum while its neutralizing binding protein (IL18BP) was significantly upregulated, suggesting that the increased IFNγ-mediated response may be driven by IL-12 without IL-18 synergy ^11,12^. Furthermore, increased expression of Th17 and the Foxp3^+^ Treg subsets in TB sputa was consistent with significantly enhanced transcription of transforming growth factor beta (TGF-β). Together, the host transcriptome in sputum shares both similarities and key differences compared to whole blood ^13^ and reveals a significant and specific anti-mycobacterial response in the airways not found in non-TB respiratory conditions.

In parallel with the inflammatory response there was a striking change in host central metabolism in TB sputa, with evidence of a switch from oxidative phosphorylation to glycolysis (**Table S3**). Expression of genes involved in the tricarboxylic acid (TCA) cycle was significantly downregulated (**Fig. 1f**) and broken after citrate, with reduced transcription of aconitase (ACO1) and elevated transcription of aconitate decarboxylase (ACOD1/IRG1) ^14^ (**Supplementary Fig. S2**). The electron transport chain (ETC) (**Fig. 1g**) was also significantly downregulated in TB sputa, including genes encoding NADH dehydrogenase, cytochrome c oxidase, ubiquinol-cytochrome c reductase and mitochondrial ATP (F_0_F_1_) synthase (**Table S3**). In contrast, there was an enhanced expression of glucose transporter GLUT1 (encoded by SLC2A1) and lactate exporter MCT4 (encoded by SLC16A3) (**Fig. 1h**), along with a significant increase in the ratio of LDHA to LDHB (lactate dehydrogenase A and B) (**Fig. 1h**) indicative of increased conversion from pyruvate to lactate ^15^. Increased transcription of genes involved in the oxidative branch of the pentose phosphate pathway (PPP) was consistent with production of NAPDH in association with generation of reactive oxygen species (ROS) (**Fig. 1i** and **Supplementary Fig. S3**), though transcripts associated with alternative NADPH-generating pathways (cytoplasmic malate dehydrogenase (MDH1), malic enzyme (ME1) and isocitrate dehydrogenase (IDH1)) were found at higher abundance in non-TB sputum. Together, these data support the notion that there is an overall reprogramming of host central metabolism during *Mtb* infection towards increased glycolysis, either as a positive feedback mechanism to maintain a fully activated immune response ^16^, or to produce glycolytic intermediates required for cell proliferation as part of antimicrobial defense ^17^.

### Microbiome landscape and its adaptation to *Mtb* infection

The inflammatory response revealed by direct transcriptional profiling of sputum samples shares key features common to responses to *Mtb* infection previously documented in cell culture models and infected human and animal tissues. We anticipated that if this transcription profile was translated into a functional antimicrobial response, it may disrupt the ecology of the commensal respiratory microbiota. To test this hypothesis, we compared overall microbiome taxonomy and the transcriptional profile of dominant commensal bacterial species between TB and non-TB sputum.

Taxonomic classification of the bacterial reads identified 30 phyla, 613 genera and 1331 species (**Table S4**). Reads mapping to sequenced bacterial genomes ranged from 10^6^ to 10^8^ and the overall taxonomic composition of our TB sputa was similar to that previously reported using 16S DNA ^18^, with *Streptococcus, Neisseria, Prevotella, Haemophilus* and *Veillonella* being the most represented genera (**Fig. 2a**). Non-TB sputa had significantly higher microbiome species richness than TB sputa (*p*<0.01 for both operational taxonomic units (OTUs) and Chao1 estimator) (**Fig. 2b**), but there was no difference in species diversity (Shannon and Simpson indices) (**Fig. 2c**), indicating that the distribution of species dominance and evenness was not affected by *Mtb* infection. In keeping with published literature, similar lung and oral microbiome diversity in HIV-uninfected and HIV-infected patients ^19^, species richness or diversity in TB sputa was unaffected by HIV-1 co-infection (**Fig. 2d**).

**Fig. 2.**
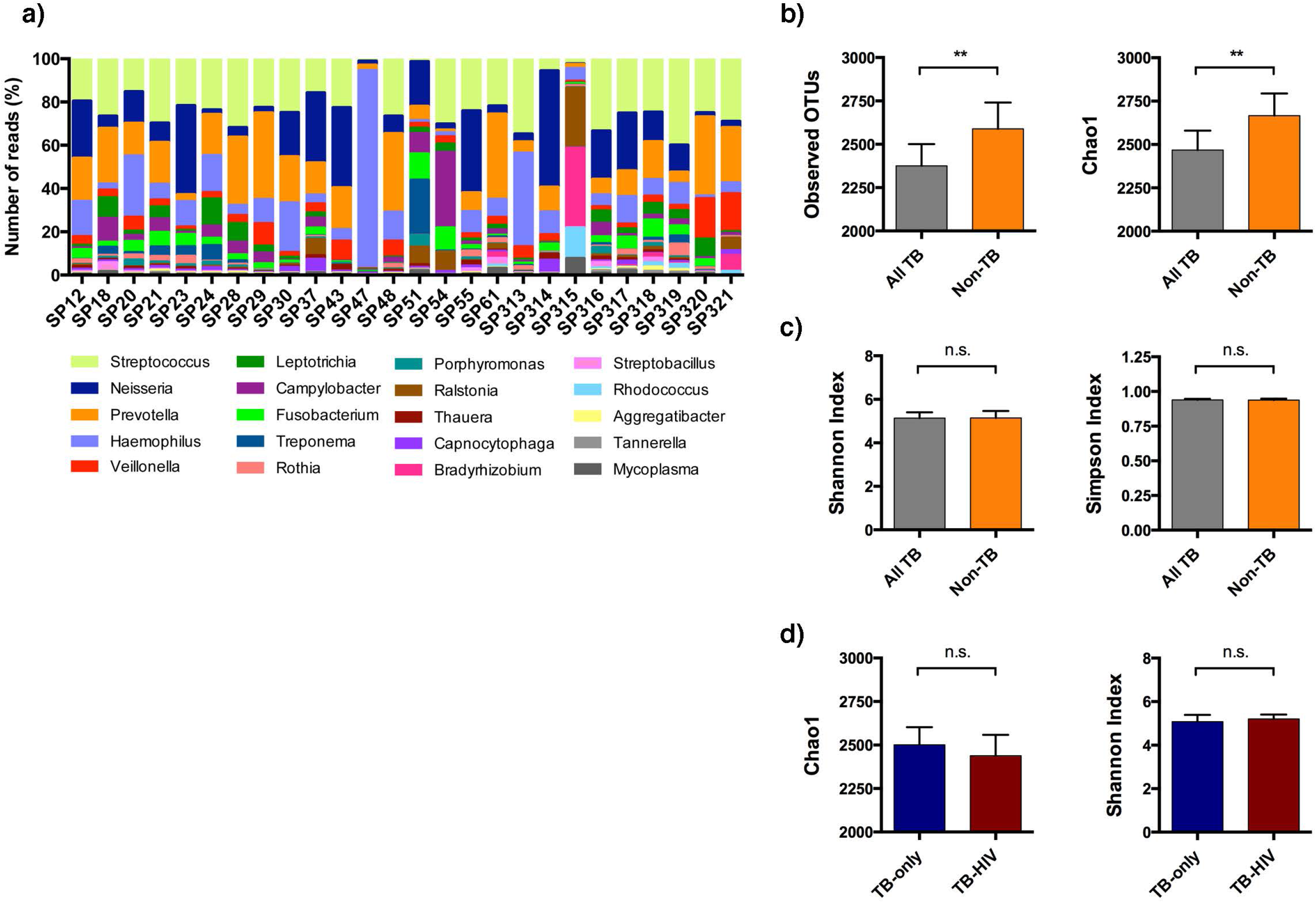
Global overview of sputum microbiome. **a)** A stacked bar chart to show the top 20 most represented microbiome genera in TB (SP12-SP61) and non-TB (SP313-SP321) sputa. SP47 had an expansion of *Haemophilus* and SP315 comprised mainly of known artefacts *Ralstonia* and *Bradyrhizobium*. These two samples were subsequently removed from all downstream analyses. **b)** Microbiome species richness and diversity were calculated. Non-TB samples (n=9) had a significantly higher number of observed operational taxonomy units (OTUs) and estimated number of true OTUs (chao1 indicator), compared to TB samples (n=17). **c)** There was no difference in species diversity as measured by the Shannon and Simpson indices, indicating species evenness and distribution did not differ between TB and non-TB groups. **d)** HIV-1 co-infection did not impact the global microbiome species richness or diversity in sputum. For panels b-d, statistical difference was calculated using Mann Whitney *U*-test and * p<0.05, ** p<0.01 and n.s. for not significant.

### Transcriptional profiling of sputum *Mtb*

Reads mapping to *Mtb* accounted for only 0.85±2% of total mapped bacterial reads (**Fig. 3a**), ranging from 10^3^ to 10^5^. Consistent with evidence of lower transmission from HIV-1 co-infected patients ^20^, there was a significantly higher percentage of *Mtb* reads in TB-only, compared to the TB-HIV sputa (mean: 1.55% *vs.* 0.06%, respectively; *p*=0.027) (**Fig. 3a**).

**Fig. 3.**
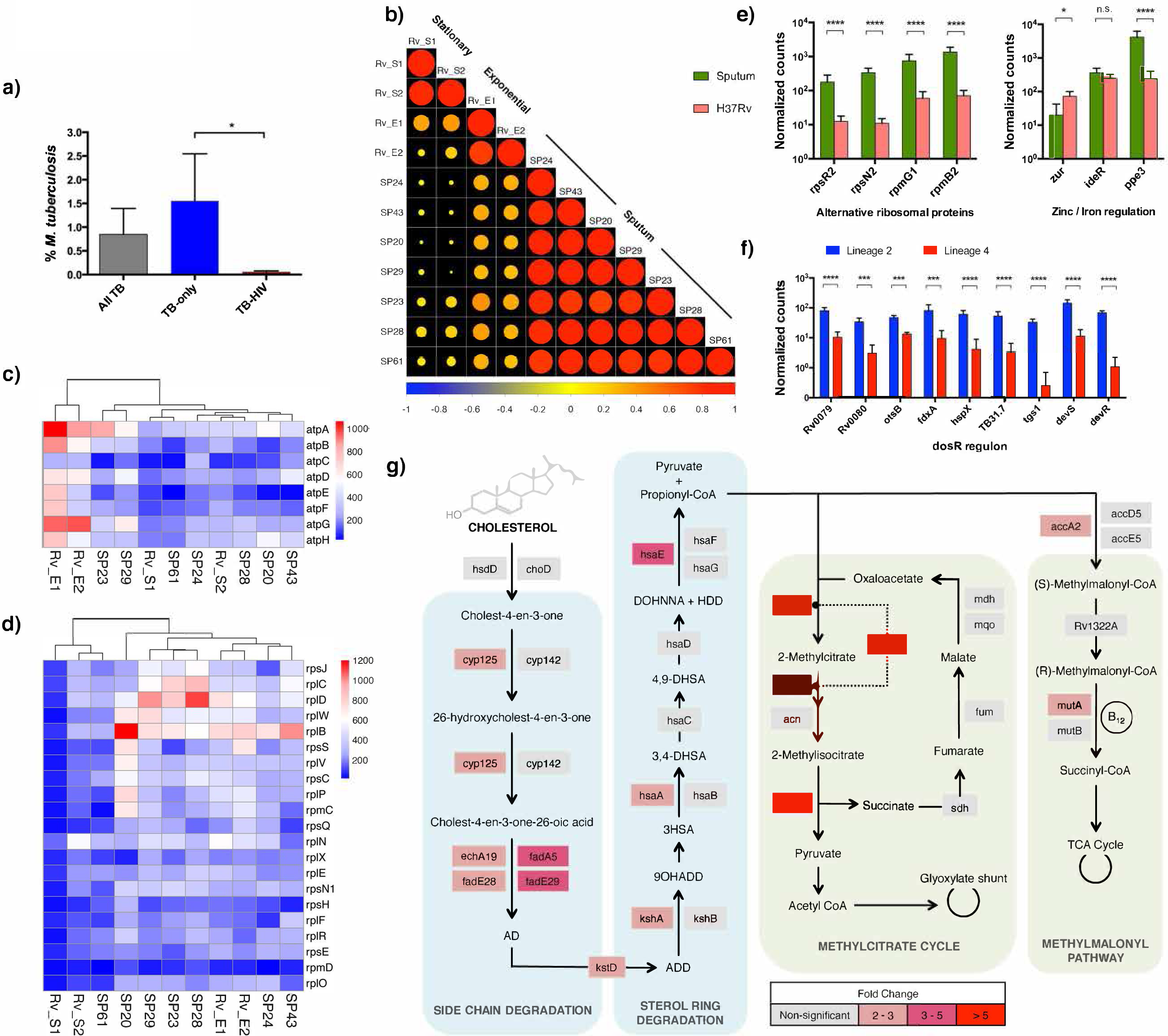
Transcriptional profiles of sputum *Mtb*. a) Despite active TB disease, *Mtb* only accounted for 0.85±2% of all mapped bacterial reads. The percentage of *Mtb* reads was, however, significantly higher in TB-only samples, compared to TB-HIV (n=9 and n=8, respectively; p<0.05, Mann Whitney *U*-test). **b)** Differential gene expression between seven sputum *Mtb* samples and laboratory cultures was calculated using DESeq2. The expression data was plotted as a correlation matrix with hierarchical clustering. Exponential cultures were labeled as Rv_E1 and Rv_E2, stationary cultures were labeled as Rv_S1 and Rv_S2, and sputum samples started with the initials SP. A decrease in circle size indicates reduced correlation; red indicates a positive correlation and blue indicates negative correlation. The sputum samples showed a high degree of concordance to each other and correlated more closely to exponential cultures than to stationary cultures. **c)** Transcript abundance of ATP synthase genes in sputum *Mtb* clusters more closely to stationary phase H37Rv than to exponential phase cultures. **d)** In contrast, transcript abundance of the two major ribosomal protein operons S10 and L14 in sputum *Mtb* were found to be more similar to exponential phase H37Rv than to stationary phase cultures. Color of the heatmaps corresponds with the normalized read count of each gene. **e)** Significantly higher expression of four zinc-independent alternative ribosomal proteins was detected, along with decreased expression of the zur repressor and upregulation of *ppe3*, indicating that sputum *Mtb* was zinc-deprived. **f)** Expression of selected members of the DosR regulon is shown. Consistent with the presence of an alternative transcriptional start sites in lineage 2 isolates ^23^, transcript abundance of the DosR genes was significantly higher in lineage 2 than in lineage 4 sputum *Mtb*. **g)** Compared to exponential phase laboratory cultures (H37Rv), *Mtb* in sputum was found to have significantly higher expression of 34 members of the KstR and KstR2 regulons associated with cholesterol catabolism and 6 members of the downstream propionate detoxification pathways. A pathway map is shown here to illustrate the transcript expression of some of the enzymes involved in the processes. Genes that were not differentially expressed (non-significant) are colored in gray color and those that were differentially expressed in sputum were colored in scale of pink and red colors according to their fold change. No downregulated genes were identified in the KstR/KstR2 regulons or either of the propionate detoxification pathways. For panels e-f, adjusted p-values (padj) were determined by DESeq2 and shown as asterisks: * padj<0.05, ** padj<0.01, *** padj <0.001 and **** padj <0.000, and n.s. for non-significant.

Seven samples (6 TB-only and 1 TB-HIV) had sufficient read coverage (>4×10^4^ reads) to quantify transcript abundance for >50% of the *Mtb* genome. Three of the samples were identified as belonging to Lineage 2, one to Lineage 3, and three to Lineage 4 (**Table S1**). In the obvious absence of a comparative control from non-TB sputa, we compared the sputum *Mtb* transcriptome to exponential and stationary phase liquid laboratory cultures of *Mtb* strain H37Rv. Plotting expression data as a correlation matrix demonstrated that the sputum profiles formed a closely related cluster that shared greater similarity to exponential than to stationary phase culture (**Fig. 3b**). Expression analysis identified 198 genes as differentially expressed between sputum and exponential culture (*p-adjusted* < 0.05; **Table S5**), and 392 genes between sputum and stationary phase (*p-adjusted* < 0.05; **Table S6**).

Transcript abundance across the ATP synthase operon in sputum was closer to stationary phase than to exponential culture (**Fig. 3c**), while transcription of the main ribosomal protein operons more closely resembled the exponential reference (**Fig. 3d**). A striking feature of the ribosomal protein gene profile in sputum was high abundance of transcripts for a set of four alternative ribosomal proteins characteristic of growth in a low zinc environment (**Fig. 3e**). Additional zinc-regulated genes ^21^ including the putative chaperone Rv0106, methyltransferase Rv2990c, and the ESX-3 operon were also significantly increased in sputum compared to laboratory culture (**Tables S5 and S6**). The ESX-3 operon is under dual control of zinc-responsive Zur and iron-responsive IdeR repressors; induction of *ppe3*, which lies upstream of the IdeR site and downstream of a Zur site, provides further indication of zinc deprivation (**Fig. 3e**). Expression of the DosR stress regulon in sputum more closely resembled the exponential than the stationary phase reference (**Tables S5 and S6**), with significantly higher expression of DosR genes in sputum samples infected with Lineage 2 compared to Lineage 4 isolates (**Fig. 3f**). Inspection of expression profiles showed that this reflected an increase in *dosR* transcripts originating from a SNP-generated constitutive start site internal to Rv3134c in Lineage 2, rather than from the stress-inducible start site upstream of Rv3134c ^22-24^ (**Supplementary Fig. S3**).

Thirty-four members of the KstR and KstR2 regulons involved in degradation of cholesterol side chain and ABCD rings ^25^, and genes involved in downstream propionate metabolism by the methylcitrate cycle ^26^ and methylmalonate pathways ^27^ were consistently higher in sputum than laboratory culture (**Fig. 3g**). This is similar to previous descriptions of the induction of *Mtb* cholesterol catabolism genes in macrophage and mouse models ^28,29^. PhoP plays an important role in transcriptional regulation during *Mtb* infection and analysis by chromatin-immunoprecipitation has identified a set of genes that are regulated by binding of PhoP to upstream sites ^30^. Twenty PhoP-regulated transcripts, including small RNA *mcr7*, were differentially expressed in sputum compared to laboratory culture; in all but one case the sputum profile was consistent with a decrease in PhoP binding (**Table S5**).

We validated 15 differentially expressed genes using NanoString methodology and compared transcript levels in three sputum samples against an independent *Mtb* H37Rv reference culture. These included representative upregulated (KstR, Zur, propionate) and downregulated (ATP and mycobactin synthesis) genes. All genes showed the same pattern of differential expression (**Table S7**) and validated the use of dual RNA-Seq in studying *Mtb* transcriptome despite its minor representation among the microbial community.

## Discussion

*Mycobacterium tuberculosis* spends most of its life sequestered in lesions within tissues, but in order to transmit to a new host it has to move into the respiratory tract prior to release in the form of infectious aerosol droplets ^31^. The transmission phase is hard to model in experimental systems and is poorly understood. We reasoned that sputum samples could be exploited to obtain additional information about conditions in the respiratory tract that may influence the efficiency of TB transmission. We generated RNA sequence data directly from sputum and analyzed these with respect to host, pathogen and microbiome transcripts to provide a comprehensive overview of the entire ecosystem. This is the first report that such strategy can be successfully applied to pathological specimens, with manifest implications for the study of other human infectious diseases to complement *in vitro* and animal models.

Comparison of host transcript profiles from TB patient sputum with *Mtb*-negative sputum revealed wholesale changes characteristic of the innate and adaptive immune inflammatory response. Given the unpromising physical appearance of sputum as a heterogeneous mixture of cell debris and mucoid secretions, the homogeneity and clarity of the transcriptional response is striking, and may reflect elimination of signal from dead cells by mRNA degradation. As in previous clinical studies using whole blood ^32^, we detected a strong type I/II interferon-mediated cytokine responses in sputum, but a strong T-cell activation and differentiation signature detected in sputum is not seen in blood; likely reflecting sequestration of these cells at the site of disease. These changes were accompanied by a metabolic shift towards glycolysis with a reduction in oxidative phosphorylation and a broken TCA cycle ^33^. The Warburg effect in mycobacterial infection is IFNγ-dependent ^34^ and probably results from a functional change in the mitochondria from energy generation to production of ROS. Upregulation of superoxide dismutase, myeloperoxidase, and glutathione peroxidase were identified in TB sputa (**Table S3**), implicating a shift in the role of host mitochondria towards bactericidal activity. A switch to glycolysis, which allows rapid production of ATP, would therefore compensate for energy loss and maintain the mitochondrial membrane potential, while upholding antimicrobial defense mechanisms.

While the majority of microbiome studies focus on the intestine, there is increasing interest in respiratory microbiota ^35^. Only a few studies have examined the microbiome in TB ^18,36,37^. The bacterial species detected by sputum RNA sequencing in our cohort are similar to those reported in other studies of the oral cavity and respiratory tract, reflecting the inevitable mixing associated with coughing and expectoration, and include a combination of aerobic and anaerobic members of firmicute, bacteroidetes and proteobacterial phyla. As reported in previous studies of the lung microbiome, we did not observe any major impact of HIV-1 infection on taxonomic distribution ^19^. We did, however, find a significant reduction in species richness in TB compared to non-TB sputum. Intriguingly, despite having active disease, *Mtb* only accounted for a very small percentage of total bacterial reads measured and was almost negatable in those with HIV-1.

We acknowledged that the total read counts detected for *Mtb* is very low for typical differential gene expression analysis. This is due to the one-step protocol in which no bacterial enrichment was performed in order to accurately assess the abundance of *Mtb* in its natural environment and to avoid induction of transcriptomic changes during the enrichment process. Despite the low read counts and its minute representation amongst total bacterial population, there was an overwhelming upregulation of genes associated with cholesterol catabolism ^29,38^. The ability of *Mtb* to utilize cholesterol is unique amongst the major species in the respiratory microbiome as *Mtb* can shunt the toxic of by-product (propionate) into the methylcitrate cycle and the methylmalonyl pathway, which may be of a crucial adaptive significance. The *Mtb* sputum transcriptome also reveals evidence of zinc deprivation. This is of particular interest in light of evidence that the bacteria face the opposite challenge of zinc intoxication when phagocytosed by activated macrophages ^39^. Neutrophil-derived calprotectin may restrict the availability of zinc in the respiratory tract, and competition with commensals for free zinc may represent a vulnerability of *Mtb* in sputum. Similarly contrasting with results in macrophage culture ^40^, the *Mtb* sputum transcriptome is characterized by reduced activation of the PhoP regulon in comparison to exponential culture. Several studies have partially characterized the transcriptome of *Mtb* from sputum or bronchoalveolar lavage using whole-genome probed-based qPCR or microarray 2,41-44. There is significant common ground in energy metabolism, ATP synthesis, iron response and PhoP regulon when comparing our data to these studies, but with key differences in the DosR regulon. Expression of DosR genes in sputum *Mtb* has been described to resemble hypoxic non-replicating laboratory cultures ^42,44^, or distinctive from both aerobic and hypoxic cultures ^2^, and found in lower abundance in HIV-1 coinfected patient samples when lineage was controlled ^45^. The discrepancies could be due to geographic location and lineage of the samples collected, sample preparation, the technology used for quantification and the growth conditions and origin of the laboratory cultures used for comparison. Finally, it will be important to determine the ratio of extracellular to intracellular *Mtb* in sputum; while there is clearly recruitment of an activated population of inflammatory cells in TB sputum, it is possible that they are engaged in phagocytosis of commensal bacteria rather than *Mtb*.

The overall aim of our research is to identify interventions that will reduce the viability of *Mtb* in the respiratory tract in order to reduce the efficiency of infection and transmission. We anticipate that this could involve vaccination to prime effective T cell responses and opsonizing antibodies, targeted antibody or small molecule therapies to optimize host responses, and nutritional or antibiotic interventions that alter the respiratory microbiome. Comprehensive mapping of the transcriptional landscape of both the host and the *Mtb* described here provides a crucial framework for further study.

## Acknowledgements

The authors thank all the participants in this study and the health care workers and administrators at the Ubuntu Clinic. We would like to thank Meena Anissi, Leena Bhaw, Deborah Jackson and Abdul Sesay of the High Throughput Sequencing facility at the Francis Crick Institute for help with the sequencing. We thank the UCL Nanostring facility for providing the nCounter system and related services. RPJL is supported by the UK Medical Research Council (MRC/R008922/1). RJW is supported by the Francis Crick Institute which receives its core funding from Cancer Research UK (FC00110218), the UK Medical Research Council (FC00110218), and the Wellcome Trust (FC00110218). Other funding included the Wellcome Trust (097254 for SM and 084323 and 104803 for RJW), European Union (FP-7-HEALTH-F3-2012-305578 for RJW and FP7 SysteMTb Collaborative Project 241587 for TC), the National Research Foundation of South Africa (96841 for RJW) and the Carnegie Corporation Training Award and Discovery Foundation Academic Fellowship Award (SM).

## Author Contributions

RPL, SM and RJW conceived and designed the experiments; SM and NR recruited, sampled and collected data from patients; RPL and MLB performed the experiments; RPL, TC, MLB, AG, SH and DBY analyzed the data; SM, NR, SH, AOG, DBY and RJW contributed materials and analysis tools; all authors contributed intellectual input; RPL, DBY and RJW wrote the paper.

