## Supplementary Figures 1-3 for "Transcriptomic characterization of tuberculous sputum reveals a host Warburg effect and microbial cholesterol catabolism"

### SI Figure S1

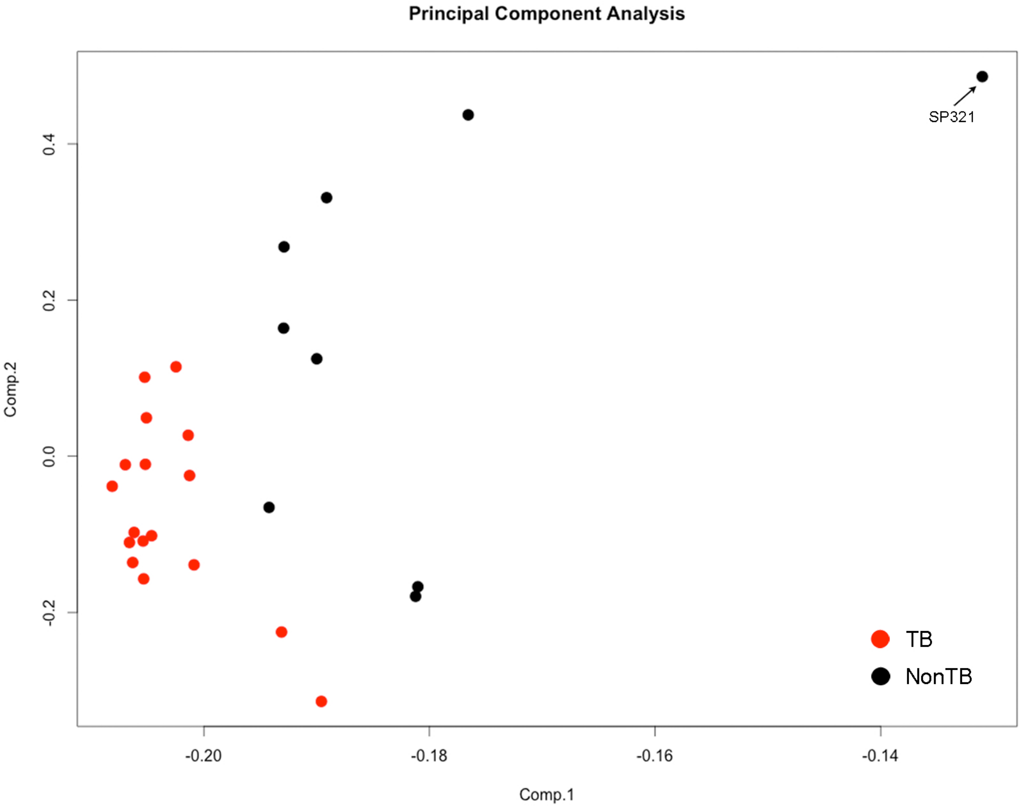


**Supplemental Data Figure S1. Principal component analysis of host transcript profiles**

The host transcriptomes of sputum samples were analyzed by principal component analysis. A complete segregation of the TB (red) from the non-TB (black) samples was observed. One of the non-TB sputa (SP321) was an outlier with differential clustering pattern and was excluded from downstream analysis of the host gene expression.

### SI Figure S2

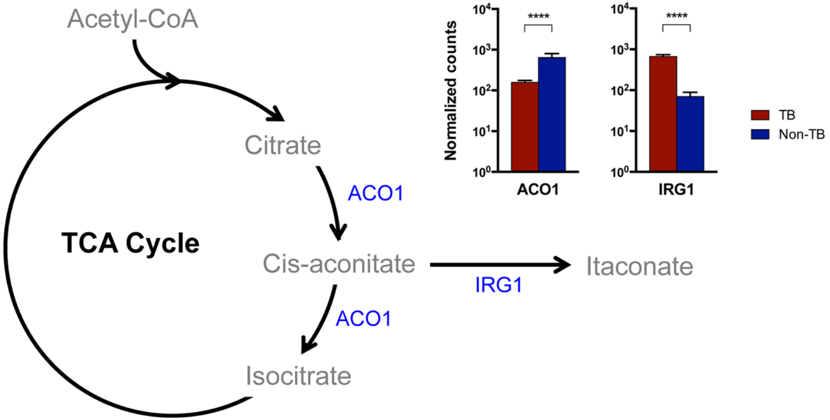


**Supplemental Data Figure S2. Itaconate biosynthesis in the host**

The TCA cycle of the host in TB sputa was similar to the pattern previously described in M1 inflammatory macrophages, broken after citrate and resulted in increased production of itaconate. The ACO1 enzyme that converts citrate to cis-aconitate and isocitrate was significantly downregulated, while IRG1 that mediates conversion to itaconate was significantly induced in TB sputa.

**SI Figure S3**

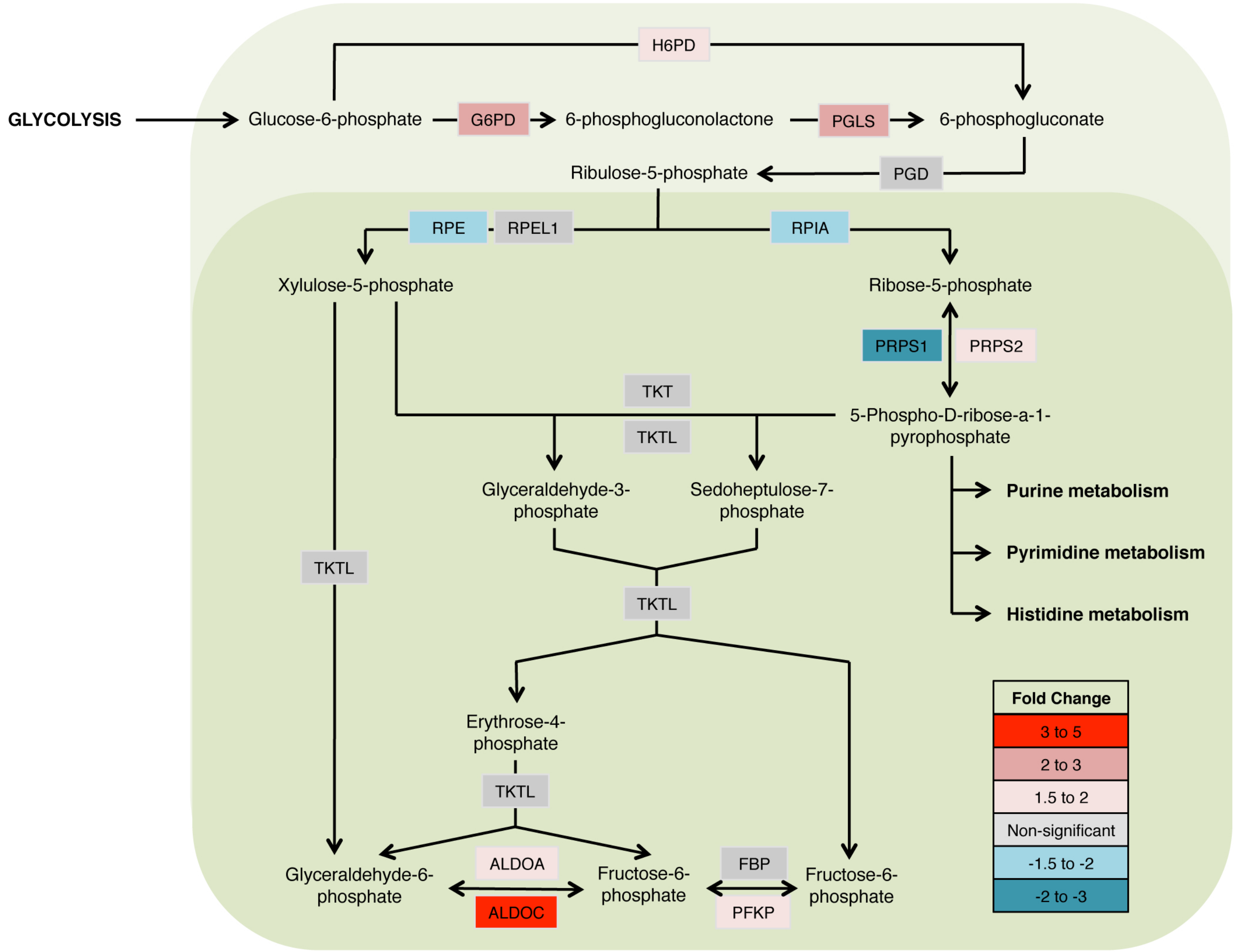


**Supplemental Data Figure S3. Pentose phosphate pathway in the host**

The pentose phosphate pathway is illustrated here. The steps in the light green shade represent the oxidative branch of the pathway involved in NADPH production. Transcript abundance of enzymes that mediate the oxidative steps was significantly higher in TB sputa compared to non-TB. In contrast, there was no change or reduction of gene expression associated with the non-oxidative branch of the pathway (shaded in dark green).
